# PII1/MRC interaction with Starch Synthase 4 (SS4) from *Arabidopsis thaliana* inhibits SS4 enzymatic activity

**DOI:** 10.1101/2024.10.02.616297

**Authors:** M. Bossu, R. Osman, G. Brysbaert, Marc F. Lensink, D. Dauvillée, C. Bompard

## Abstract

Starch is the major energy storage compound in plants. It accumulates in the form of insoluble, partly crystalline granules whose number and shape are specific to each plant species. These characteristics are defined very early in starch biosynthesis, at the initiation stage.

Starch biosynthesis initiation is a complex process that relies on the coordinated action of several proteins that interact together in the so-called complex of initiation. Starch Synthase 4 (SS4) is the only initiation protein with enzymatic activity. It catalyzes the formation of glucan primers, which serve as substrates for the enzymatic machinery that synthesizes starch granules. Previous studies have highlighted the importance of interactions between SS4 and regulatory proteins in this process. Among them, Protein Involved in Initiation 1 (PII1) interacts with SS4 but its function is not yet established. In this study, we explored the structural and functional implications of PII1 on SS4’s enzymatic activity. Our findings reveal that PII1 contains a long coiled-coil domain that specifically interacts with SS4, leading to significant inhibition of SS4’s glucan elongation activity. Importantly, this inhibition is specific to SS4 and does not affect other known synthases, suggesting a targeted regulatory mechanism. This work describes the structural specificities of PII1 and SS4 and reveals a function for PII1 in the initiation complex. These results allow us to re-examine these complex mechanisms and propose new hypotheses about the important steps in the initiation of starch biosynthesis.

## Introduction

In higher plants, starch is the most abundant and widely distributed non-structural carbohydrate that accumulates as water-insoluble, partly crystalline granules. In heterotrophic tissues, starch functions as a long-term carbohydrate reserve, supporting germination or seasonal regrowth. This “storage starch” is a key component of our staple crops, accounting for half of human caloric intake and is also extensively used as an industrial commodity. In leaves, transitory starch accumulates in chloroplasts during the day and is used as carbon and energy source during the night.

Starch is composed of two polymers of glucose residues, namely amylose and amylopectin, organized as linear α-1,4-glucans covalently linked to one another by α-1,6-bonds also called branching points (for review (Pfister and Zeeman, 2016)). Amylose molecules are mainly linear (<1% α-1,6 bonds) whereas amylopectin, the main component contains 5-6% of branching points (Pérez and Bertoft, 2010; Pfister and Zeeman, 2016).

Amylose linear chains are synthesized by the action of the Granule Bound Starch Synthase (GBSS) that is embedded in starch granules and belongs to the starch synthase class of glucosyltransferases (Ball and Morell, 2003; Schwarte et al., 2013; Pfister and Zeeman, 2016; Seung, 2020). A second non enzymatic protein, namely PROTEIN TARGETING TO STARCH 1 (PTST1) is also involved in the process by targeting GBSS to starch granules (Seung et al., 2015).

Amylopectin is synthesized by the concerted activities of soluble starch synthases (SSs), starch branching enzymes (SBEs) and starch debranching enzymes (SDBEs) acting by forming transient complexes during the different stages of biosynthesis (Crofts et al., 2015). Soluble SSs transfer the glucose residue of ADP-Glucose (ADP-glc), the precursor molecule, to the non-reducing end of growing amylopectin and amylose molecules, or on free malto-oligosaccharides (MOSs) that arise from degradation of starch granules formation (Larson et al., 2016; Seung et al., 2017; Xie et al., 2018).

Branching points are introduced by SBEs which cleave and transfer a linear malto-oligosaccharide and transfer it to an α-1,6 position (Sawada et al., 2014). The isoamylase class (ISA) of SDBEs is involved in the synthesis of amylopectin by hydrolyzing the excess and incorrectly positioned α-1,6 bonds of soluble amylopectin molecule thus facilitating amylopectin crystallization (Ball et al., 1996; Myers et al., 2000; Delatte et al., 2005; Wattebled et al., 2005). Finally, two recently discovered non enzymatic proteins, LESV (LIKE EARLY STARAVTION 1) and ESV1 (EARLY STARVATION 1) are involved in amylopectin packing and starch granule protection (Liu et al., 2023; Osman et al., 2024).

In addition to these biosynthetic mechanisms, an initiation step is required to produce the oligosaccharide substrates of the biosynthetic enzymes. This step has been identified, but its molecular mechanism remains poorly understood even if several proteins involved in this mechanism have been identified. The first of them is an enzyme from the SSs family, starch synthase 4 (SS4). Arabidopsis mutant plants lacking SS4 show a phenotype in which the number of starch granules formed in the chloroplasts is greatly reduced and the remaining granules are much larger and spherical, in contrast to the lenticular granules observed in the wild-type plants (Roldan et al., 2007). This 1040 amino acids protein is composed of three distinct domains: an N-terminal domain containing a transit peptide (1-42), a predicted coiled-coil region (43-465), a so-called dimerization domain (471-515), and the catalytic domain at the C-terminus (540-1040). The N-terminal domain is predicted as a coiled-coil (cc) domain containing four cc regions (Raynaud et al., 2016). This N-terminal domain was shown to be required for the granule shape, while the catalytic domain is sufficient to regulate granule number (Lu et al., 2018). Subsequently, several non-enzymatic proteins were identified as being involved in starch granule initiation. Arabidopsis plants that do not express these proteins show a phenotype in which the number of granules is reduced, but they keep their lenticular shapes (Lu et al., 2018). These proteins include two members of the PTST family: PTST2 and PTST3, two conserved coiled-coil and CBM48 containing proteins (Seung et al., 2017), the thylakoid-associated MAR-BINDING FILAMENT-LIKE PROTEIN 1 (MFP1) (Sharma et al., 2024), MYOSIN-RESEMBLING CHLOROPLAST PROTEIN or PROTEIN

INVOLVED IN INITIATION 1 (MRC/PII1)(Seung et al., 2018; Vandromme et al., 2019) and a non-canonical starch synthase, SS5 (Abt et al., 2020).

It has been proposed and shown that these proteins interact specifically, probably sequentially and in some cases transiently, as part of an initiation complex whose molecular mechanism is still poorly understood (Seung et al., 2018). The function of some of these proteins has been investigated.

The first event in starch initiation is likely linked to the action of the MFP1 that interacts with PTST2 and specifically determines the subchloroplast location of the starch granule initiation machinery (Sharma et al., 2024). PTST2 also interacts with PII1 and SS4 but interaction with SS4 is not dependent of the cc region (Seung et al., 2017). Through its CBM, it binds and delivers glucans, able to adopt a helical secondary structure, to SS4. The function of PTST3 is partially redundant with that of PTST2 (Seung et al., 2017). The non-canonical starch synthase, SS5, which contains an incomplete catalytic domain, has no enzymatic activity and interacts with PII1 (Abt et al., 2020). It remains to be determined whether proteins with multiple partners are able to interact with them simultaneously *via* multiple recognition sites, or whether these interactions are in competition with each other as part of their functional regulation.

PII1 has a predicted coiled-coil region that spans most of the protein without any predicted catalytic or glucan binding domains (Seung et al., 2018). Knockout mutants have a phenotype analog to *ss4* mutant plants with one large lenticular (rather than spherical in *ss4*) granule per chloroplast instead of the normal 5-7. This phenotype is similar to the *ptst2* phenotype, but is less severe, as *pii1* chloroplasts still contain one starch granule in most chloroplasts, whereas *ptst2* and *ss4* mutants have many empty chloroplasts. PII1 function is unknown, particularly the role of its interaction with SS4 in the initiation process. In this work we studied both the structure of PII1 and SS4 as well as their interaction. Through biochemical experiments using the recombinant proteins, we were able to reveal an inhibitory effect of PII1 on the elongation capacity of SS4. This unexpected result is discussed in terms of its potential implication on the regulation of starch granule initiation *in vivo*.

## Methods

### Molecular Modelling

Protein structures of all PII1 and SS4 constructs were initially modelled using both AlphaFold2 combined through MassiveFold (ref here: https://www.researchsquare.com/article/rs-4319486/v1) that allows to go further by generating a large number of molecular models, all the neural network model versions available to date (v1, v2 and v3) (Jumper and Hassabis, 2022). Initially, three predictions per neural network model were generated for monomers, resulting in fifteen predictions per monomeric protein, and five predictions per neural network model for protein complexes, resulting in seventy-five predictions for each. Massive sampling and increased recycling were also attempted for some complexes, pushing predictions until six hundred, but didn’t show substantially improved results in these cases, compared to a basic prediction run.

When this was available, we used AlphaFold3 for the calculation of new molecular models and compared them to the predictions already obtained by MassiveFold. The structure models obtained with both approaches being similar for our proteins, we only kept AlphaFold3 predictions for the sake of clarity of the manuscript.

With AlphaFold3, for each monomeric protein, five different predictions were computed and ranked by global pLDDT. The five molecular models generated have been superimposed and used to identify the presence of dynamic regions. Molecular models with best pLDDT and PAE values were used for figures. For each complex, fifty predictions were computed and ranked by the AlphaFold confidence score. Scores and PAE values were used to evaluate the quality of the predictions.

Structures and electronic surfaces were visualized using the PyMOL Molecular Graphic System, Shrödinger, LLC.

### Synchrotron radiation circular dichroism

Synchrotron radiation circular dichroism (SR-CD) spectra were measured at the DISCO beamline of the SOLEIL Synchrotron (Gif-sur-Yvette, France). The beam size of 4 × 4 mm and the photon-flux per nm step of 2 × 10^10^ photons s^−1^ in the spectral band from 270–170 nm prevented radiation-induced damage (Miles et al., 2008). CD spectra were acquired using IGOR software (WaveMetrics). Before measurements the molar elliptical extinction coefficient of Ammonium *d*-10-Camphorsulfonate Ammonium (CSA) has been measured on the beamline and used as standard for calibration of all data measurements (Miles et al., 2004). Protein and buffer spectra were collected consecutively and are the mean of 3 accumulations. The buffer baseline was then subtracted from the spectra and the data processing was conducted using CDToolX software (Miles and Wallace, 2018).

PII1-H2-H3 protein solutions were deposited between 2 CaF_2_ coverslips with a pathlength of 10 (TFE) or 50 (protein alone) µm (Refregiers et al., 2012). The influence of the different concentration of TFE on the structure of PII1 was studied by mixing the protein and different concentrations of TFE and then measuring the spectra under the same conditions as for the native protein after a 10 min incubation. Spectra containing protein buffer at corresponding TFE concentrations were subtracted from the protein/TFE spectra before CSA calibration.

### Cloning, expression and purification of proteins

PII1-H2-H3 and SS4-Δ349 were cloned and expressed in *Escherichia coli* as recombinant proteins lacking their N-terminal transit peptides. The complete cDNAs encoding the complete PII1 and SS4 lacking their N-terminal transit peptides, cloned into the pENT-D-Topo plasmid (Thermofisher, Rochester, USA)(Vandromme et al., 2019) were used as templates for the construction of the vectors expressing the different truncated forms of SS4 and PII1. A fragment of the *Pii1* cDNA corresponding to the H2 and H3 helixes was amplified with primers EntPII1For et EntPII1Rev using the ultra-high Kapa Hifi Hot start polymerase (Roche diagnostics, Austria) and the 1.4kb PCR product was transferred into the pENT-D-Topo plasmid (Thermofisher, Rochester, USA). This entry vector was used to construct the pET300-PII1 vector by using the gateway technology into the Champion^TM^ pET300 plasmid (Thermofisher) that allows the generation of recombinant proteins fused to a N-terminal 6-Histidine tag. The PCR product corresponding to the SS4-Δ349 truncated *ss4* cDNA fragment was amplified using the Kapa Hifi Hot start polymerase and digested by *Bam*HI and *Xho*I restriction enzymes before its cloning into the pET-DUET-1 (Novagen) vector’s corresponding restriction sites allowing the expression of N-terminal His-tagged proteins. For co-expression experiments, the *PII1* sequence was amplified using the BamPII and SalPII primers allowing the introduction of *Bam*HI and *Sal*I restriction sites which were used to clone the digested PCR product into the pET-DUET-1. A second PCR product corresponding to the SS4-Δ349 cDNA sequence was amplified using the RVSS4 and XhoSS4 primers and the PCR product was cloned in the *Eco*RV and *Xho*I restriction sites of the previous plasmid allowing co-expression of both N-terminal His-tagged PII1 and SS4-Δ349 proteins.

Primers sequences are detailed in table 1, the restriction enzyme sites used for the cloning are underlined in the primers’ sequences. Each construct was fully sequenced before use to ascertain the integrity of the cloned sequences.

**Table 1:**
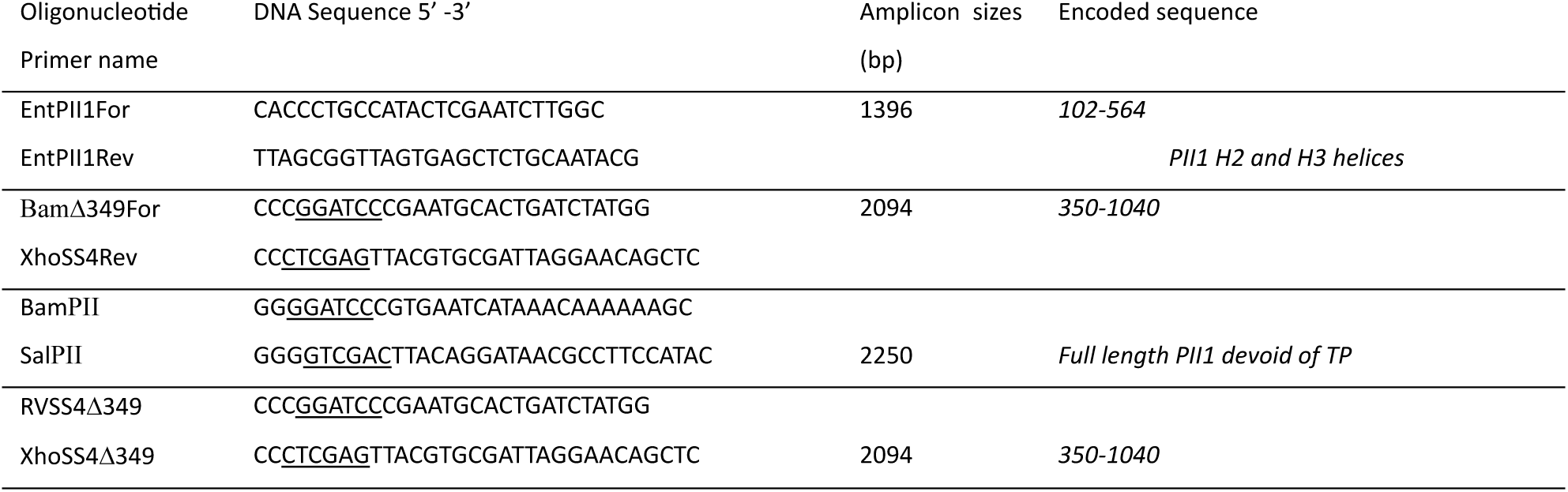
List of oligonucleotide primers used in this study. The restriction enzyme sites are underlined.

The recombinant proteins were expressed in *E. coli* BL21 DE3 in 2 x 500 ml of LB medium supplemented with 100μg/mL antibiotic then induced with 0.5 mM IPTG overnight at 20°C. Cells were pelleted at 6000 g for 30 min at 4°C and stored at -80°C for further use.

Cells pellets were resuspended in lysis buffer (20 mM Hepes/NaOH pH 7, 150 mM NaCl supplemented with one tablet per liter of EDTA-free of protease inhibitors EDTA free (Roche), disrupted by Emulsiflex or sonication and centrifuged at 10000 g for 30 min at 4°C. The supernatants were then subjected to purification.

Purification was performed by a first step of Immobilized Metal Affinity Chromatography (IMAC) using a 5ml IMAC HisTrap Excel column (Cytiva, Amersham,UK). Protein sample was loaded onto the column pre-equilibrated with buffer A (20 mM Hepes/NaOH pH 7, 150 mM NaCl). The beads were washed with 10 column volume of buffer A supplemented with 10mM Imidazole and eluted with buffer A supplemented with 500 mM Imidazole and 10% (w/v) glycerol.

This initial step was followed by a second purification step through size exclusion chromatography Superdex200 10/300 (Cytiva) equilibrated with buffer A. 500µl of the IMAC eluted sample were injected on the column. The protein sample purity was assessed by SDS-PAGE 10%.

For structural study, protein samples were concentrated using Vivaspin centrifugal concentrator with a 10 kDa cut-off (Sartorius). Protein concentrations were determined using a Nanodrop Spectrophotometer (ND1000) from Thermo Scientific.

### Zymogram

To study the elongation capacity of SS4, soluble proteins were separated by electrophoresis onto a 10 % non-denaturing acrylamide gel containing 0.3 % oyster glycogen (Sigma). The run was carried out for 2 hours at 200 mA at 4°C using the Bio-Rad Mini Protean III system. The gel was then incubated overnight at room temperature in synthase incubation mix (67 mM Glycyl-glycine/NaOH pH 9; 133 mM ammonium sulfate; 80 mM MgCl_2_; 0.6 mg.ml^-1^ bovine serum albumin; 25 mM β-mercaptoethanol; 2.4 mM ADP-glucose). Control experiments were carried out following the same procedure but omitting the ADP-glc in the incubation mix. To test the capacity of SS4 to synthesize *de novo* glucan primers, the same procedure was used but without providing any polysaccharide substrate in the gel matrix. The initiation capacity of the enzyme was tested in a control experiment in which a mix of small MOS were provided in the incubation mix.

## Results

### Expression of PII1 and SS4

To investigate the role of PII1 in the initiation steps leading to new starch granules, we aimed to study its structure as well as its interaction with SS4. For these studies, we considered a biochemical and structural approach. As these approaches required the expression and purification of both proteins, we undertook the expression of both proteins in their entirety, with the exception of their chloroplast transit peptides (cTPs). To do this, we carried out a series of expression assays on plasmids containing the genes encoding each protein, optimized for expression in *E. coli* (see Methods section). We succeeded in expressing PII1 under these conditions, but none of the tests allowed us to obtain enough soluble protein, while SS4 was little or no expressed. At this stage, and in order to continue the study, we considered using truncated but functional constructs of each protein. To identify the regions that might affect expression or solubility, we analyzed the sequence of PII1 and SS4 and calculated molecular models using AlphaFold3.

### Structural analysis of PII1 molecular model

Previous sequence analysis of PII1 predicted the presence of coiled-coil regions along almost the entire length of the protein (Seung et al., 2018; Vandromme et al., 2019) suggesting that the protein is involved in interactions with one or more other proteins. To visualize their arrangement, we then analyzed molecular models of the protein using AlphaFold3 (Abramson et al., 2024).

AlphaFold 3 computes five predictions and sort them by confidence score. Several criteria for the validity of the structure are available, the pLDDTs, the PAE matrix and, for the multimers, the TM coefficients. The pLDDTs indicates the confidence estimate per atom on a scale of 0-100, where a higher value indicates a higher confidence for each protein amino acid. The PAE (Predicted Aligned Error) matrix estimates the error in the relative position and orientation between two parts in the predicted structure. Higher values indicate higher predicted error and therefore lower confidence. And then, TM scores (predicted template modeling (pTM) score and the interface predicted template modeling (ipTM) score) that are both derived from a measurement called the template modeling (TM) score. pTM is an integrated measure of how well AlphaFold-Multimer has predicted the overall structure of the complex. In contrast, ipTM measures the accuracy of the predicted relative positions of the subunits forming the protein-protein complex. Disordered regions and regions with low pLDDT score may negatively impact the ipTM score even if the structure of the complex is predicted correctly. Both measure the accuracy of the entire structure (Zhang and Skolnick, 2004; Xu and Zhang, 2010), for a high confidence it should be >0.5 and >0.8 respectively. TM scores should be interpreted cautiously (Abramson et al., 2024) as they can be influenced by several factors. For this study all TM scores were below the “trustable” values and this can be attributed to the presence of large cc regions as well as predicted unfolded regions in proteins. Indeed, in cc regions, local contacts can be inferred with a good score, which explains the pLDDT values > 0.80 in these regions. However, the contacts can be shifted between repeats in the prediction, which makes the process less straightforward, resulting in low pTM scores. Disordered regions are always hard to predict, with high flexibility, resulting in low pLDDT values in these regions and decreasing the pTM scores. Therefore, we remained cautious in interpreting the results and models in which the PAE matrix showed no credible interactions between proteins were considered invalid.

For the full length PII1, the five models calculated by AlphaFold3 were very similar. For the sake of clarity, only the model with the best score is presented. The model obtained is composed of 6 helices predicted by a majority with a plDDT>90 confidence index and numbered H1 to H6 from the N-terminus. The 70 amino acids at the N-terminus and the region between helices H3 and H4 are predicted to be disordered. Helices H2 and H3 are very long helices of 203 and 245 amino acids respectively, organized in antiparallel coils as suggested by the sequence analysis (Figure 1A). The analysis of the PAE (Figure1B) matrix shows that the relative position of helices H1, H2 and H3 is predicted with a high confidence index, which is not the case for the other helices within them, with the exception of short helices H5 and H6.

**Figure 1:**
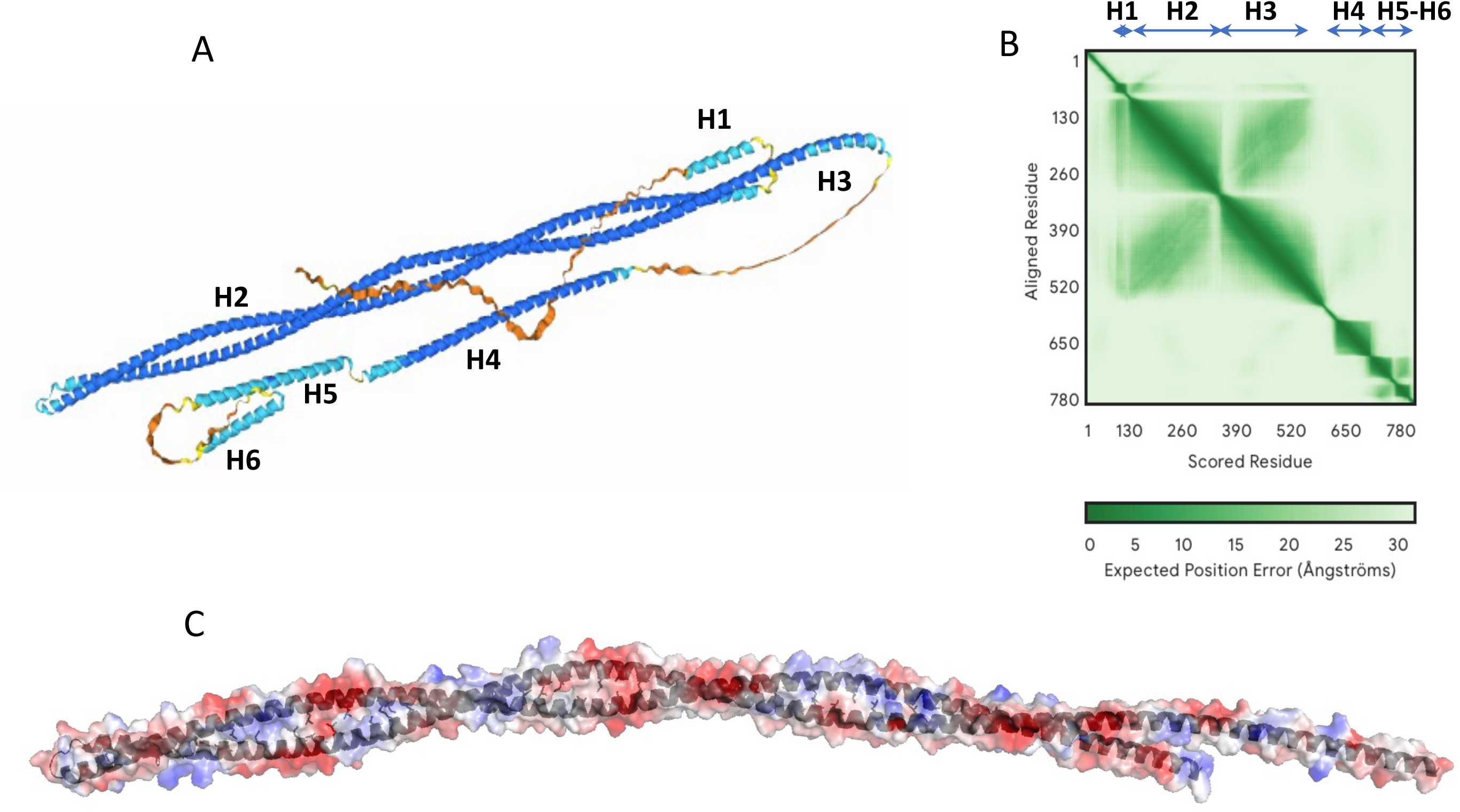
Molecular structure of PII1. The structures are represented in cartoon A) AlphaFold3 model of PII1, regions with pLDDT > 90 are colored navy blue, regions with 90 >pLDDT > 70 are colored cyan, regions with pLDDT < 70 are colored yellow and orange. The six helices of the model are labeled H1 to H6 from the N-terminal end of the protein. B) PAE matrix of the model, the region corresponding to the six helices are labeled. C) potential surface of the coiled-coil region (H2-h3) of PII1, the positive and negative charges at the surface are colored blue and red respectively. The hydrophobic side chains of amino acids involved in the coiled-coil stabilization are shown in black sticks.

The H2 and H3 helices interact with each other *via* hydrophobic interactions involving amino acids predicted to be involved in predicted repeated heptads for coiled-coils (Figure 1C). On the other hand, numerous charged residues are directed towards the protein surface and may be involved in interactions with PII1 partners. The numerous hydrophilic residues present on the surface of the H2-H3 domain of PII1, may create interactions with the solvent, increasing the chances of obtaining a soluble protein sample for its structural and functional analysis (Figure 1C).

### The coiled-coil region of PII1 is conserved among orthologs

To verify the conservation of the coiled-coil (abbreviated cc in the rest of the manuscript) H2-H3 region of PII1, we investigated whether it was conserved in its orthologs. Twenty-eight orthologs were identified on the Arabidopsis information resource (TAIR https://www.arabidopsis.org). All the orthologs are proteins close in size to *At*PII1 (between 720 and 820 residues), with the exception of the *Setaria italica* protein, which has a much shorter sequence (492 residues). Their amino acid sequences were aligned using blastp (Altschul et al., 1990) (Figure 2A, supplementary Figure 1).

**Figure 2:**
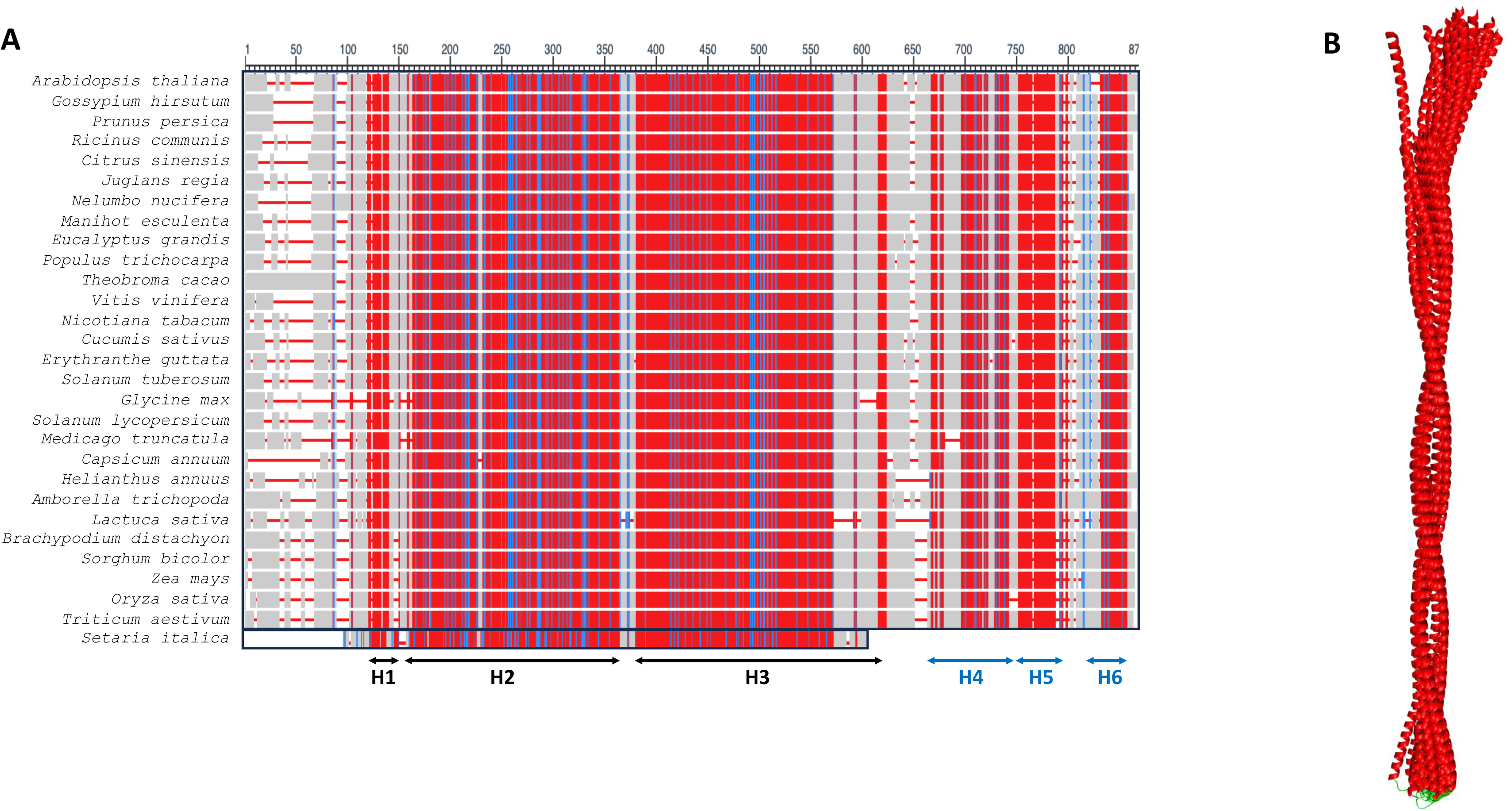
Primary and tertiary structure alignment of AtPII1 with 28 orthologs. A) Amino acid sequence alignment of PII1 with the 27 orthologs having the same length computed with Blastp. The sequence of *Setaria italica* has been manually added from the alignment of PII1 with all tested orthologs. Both alignments are detailed in Supplementary Figure 1. Only alignment positions with no gaps are colored. Red indicates highly conserved positions and blue indicates lower conservation, gaps are represented by red lines and grey regions correspond to non-conserved regions. The position of conserved helices is indicated by arrows and labeled below B) Structure alignment of H2-H3 coiled-coil region of PII1 with the 28 orthologs. Structures are represented in cartoon, helices and loops are colored red and green respectively.

The result shows high conserved sequence for the 27 *At*PII1 orthologs proteins analyzed, all along the sequence. *Setaria italica* has a much shorter sequence than the other orthologs, corresponding only to the N-terminal region containing H1, H2 and H3 in *At*PII1 (Figure 2A supplementary Figure 1).

As with *At*PII1, we calculated a molecular model for each of the identified PII1 orthologs using AlphaFold3. With the exception of the *Setaria* ortholog, which contains only the H1 and the coiled-coil domain, analysis of the results shows the presence of the 6 predicted helices similar to those predicted for the Arabidopsis protein (Figure 2A).

Alignment of the structures of H2 and H3 of PII1 and those of its 28 orthologs shows a high degree of structural conservation of the conserved cc region (Figure 2B), which is, in agreement with an important role for this region in the function of PII1.

### Dimerization domain has a crucial role in SS4 dimer structure

The structure of the catalytic domain of *At*SS4 has been solved by crystallography in the presence of ADP-glucose and acarbose, a competitive inhibitor used in the treatment of certain types of diabetes. Analysis of the structure revealed a monomeric protein and has allowed the precise characterization of its catalytic domain (Nielsen et al., 2018).

Four cc regions (cc1 to cc4 numbered from the N-terminus) have been identified in the N-terminal part of SS4 (Raynaud et al., 2016) followed by a so-called dimerization domain upstream the catalytic domain. In this work, SS4 has been shown to be dimeric and that it is in this form that it is the most active. The dimerization is linked to the presence of the dimerization domain, and cc regions are not involved in. The cc1 and cc2 regions are required for SS4 localization to thylakoid. Interestingly, the authors produced a truncated construct of the cc1 and cc2 regions (comprising the first 349 amino acids of the protein) which proved to be soluble (Raynaud et al., 2016) and, in order to obtained soluble protein samples, we decided to work with the same construct, which we will call SS4-Δ349, for the remainder of this work.

The dimerization domain is predicted to be located between amino acids 471 and 515 and its structure is not known (Raynaud et al., 2016). In order to identify this domain, to verify the possibility of dimerization and to visualize the position of the different regions that make up SS4, we created dimer models of SS4 and SS4-Δ349 using AlphaFold3.

The two models obtained show the regions containing the predicted catalytic and dimerization domains with a high level of confidence (pLDDT above 90 and between 70 and 90 respectively) and the analysis of the PAE matrix shows a good level of confidence in the relative positions of these domains (Supplementary Figure 2A, B, C, D). The AlphaFold3 predicted catalytic domain of SS4 is almost identical of the X-ray-crystallographic structure (Nielsen et al., 2018) with a rmsd of 0.3 Å for all Cα. On the predicted structure, SS4 has a predicted dimeric structure in which the dimerization domains of each monomer interact with the GT5 subunit of the catalytic domain of the second monomer sharing a large surface area (Figure 3A). The pattern of the dimerization domain, originally predicted to be between residues 471 and 515, is composed of residues 466 to 533 in the model. Its structure is predicted as a globular domain composed of a bundle of 4 helices (dH1 to dH4) parallel to each other and an N-terminal region of about 7 amino acids forming a short helix (Figure 3B). The two dimerization domains are organized into swap domains, each of which interacts with the GT5 domain of the neighboring molecule (Figure 3A). This interaction involves the helices dH1 and dH2 of the dimerization domain and the GT5 domain of the adjacent molecule described in Figure3 B, D. The interaction between the dimerization domain and the GT5 domain involves numerous hydrophilic and hydrophobic interactions between the side and main chains of residues of about ten amino acids on each side. The interaction zone covers a large surface area of both domains, indicating a strong interaction. This interaction between dimerization domain and GT5 is conserved in complete or truncated structure of SS4 (Figure3) which is a further vote of confidence in the already very convincing pLDDT and PAE matrix values (Supplementary Figure 2 C, D). In the SS4 full length model, interactions are predicted between the two dimerization domains by the short N-terminal helix (supplementary Figure 2E). In the truncated model SS4-Δ349, the interaction between the dimerization domains and the GT5 domains is identical to the full SS4 model (rmsd 0,144 Å for all Cα), with the same confidence indices and positioning errors of the same order (Supplementary Figure2C, Figure 3D). In contrast, the interaction between the dimerization domains observed in the SS4 is not conserved in the model of SS4-Δ349 in which the two dimerization domains are not in the same position and are not directly interacting (supplementary Figure 2E, F). This difference in the predictions is unlikely due to the full or truncated version of the dimer to predict, but rather to the uncertainty in the positioning of the dimerization domains between each other, as shown in both PAE matrix plots. This could indicate that, in the context of conformational changes associated either with the binding/release of glucans or with interaction with another protein in the initiation complex, the interaction between the dimerization domain of one monomer and the GT5 domain of the adjacent monomer would remain stable, whereas the interaction between the two dimerization domains could change.

**Figure 3:**
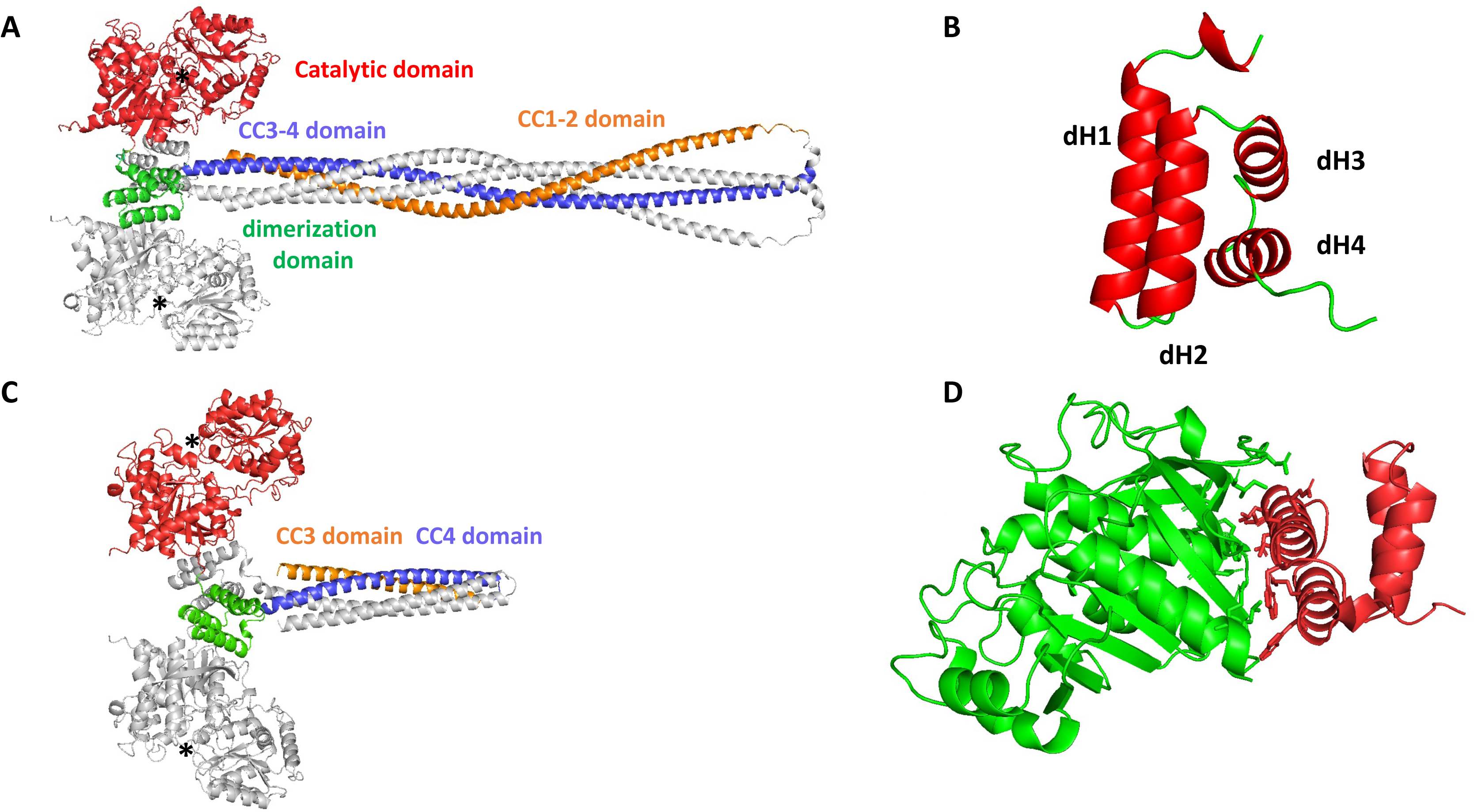
Molecular models of dimeric SS4 and SS4-Δ349. The structures are represented in cartoon A) structural organization of SS4 full length as predicted by AlphaFold 3. Only one monomer is colored (catalytic domain in red, dimerization domain in green, cc3-4 helix in violet and cc1-2 helix in orange). The catalytic sites are indicated by a black star. The second monomer is colored gray. B) molecular model of the dimerization domain of SS4. The positions of helices are labeled dH1 to dH4 from the N-terminal. C) structural organization of SS4-Δ349 with the same color code as SS4 D) Interaction between the dimerization (in red) and GT5 (in green) domains of SS4. The side chains of amino-acids residues involved in the interaction are represented in sticks.

### N-terminal domain of SS4 organizes in coiled-coil

The N-terminal domain, although predicted as helices for the cc1 to cc4 regions, have different configurations depending on whether the whole protein or the truncated protein is considered (and on the position of the dimeric domains in relation to each other).

In the five SS4 models generated by AlphaFold3 the structure of the catalytic and dimerization domains is conserved in contrast to the N-terminal domain. The latter is always predicted to be organized as a coiled-coil, but the distribution of helices, predicted with a lower confidence index than the C-terminal domain, is quite different. In the best model, as expected, the 1-180 region is predicted to be disordered, while the cc1/cc2 and cc3/cc4 regions are involved in two distinct 200 Å long helices. The cc3/cc4 helices of each monomer are predicted with good confidence (pLDDT between 90 and 70) whereas the cc1/cc2 helices are predicted with low confidence (pLDDT <70) (supplementary Figure2 A to D). Altogether, in the model they form a tetrameric coiled-coil confirmed by the PAE. In this organization, the two cc3/cc4 regions form a central dimeric coiled-coil around which the cc1 and cc2 regions are supercoiled (supplementary Figure3 A). It seems likely that AlphaFold positioned cc1 and cc2 on cc3/cc4 to stabilize the latter. cc1 and cc2 have been shown to be involved in (and essential for) the localization of SS4 to thylakoids. In this case, they must form a domain distinct from cc3/cc4 in order to interact with MFP1 or PTST2. It is therefore highly likely that they are not organized as in the model described.

The five SS4-Δ349 AlphaFold models were also very similar. In these models the cc3 and cc4 regions form distinct helices forming an antiparallel cc that assembles with the cc3/cc4 region of the second molecule, whose folding is identically predicted, to form a confident tetrameric cc (90> pLDDT >70 and 70> pLDDT >50) folded with low expected position error (supplementary Figure 2C, D, supplementary Figure 3B). This configuration is probably adopted to stabilize the hydrophobic regions of cc3/cc4 that interact with cc1/cc2 in the model of the whole protein. Like for the full-length protein, this could indicate that individual cc regions are able to adapt their folding depending on the partner with which they interact. This would give SS4 the ability to adapt to multiple partners (individually or together) by adjusting the conformation of its cc regions.

### SS5 has a similar dimeric organization as SS4

Since SS5 is a protein involved in the initiation mechanisms and interacts with PII1, we studied its structure using AlphaFold3 (Abramson et al., 2024). SS5, like SS4, is predicted to be dimeric (Abt et al., 2020), so we started by modelling a dimer. The model obtained has a high confidence index, as shown by the pLDDTs per residue and the PAE matrix (supplementary Figure 4A, B).

The first 59 amino acids are predicted to be disordered and residues 60-139 of the two molecules form two 115 Å long helices forming a coiled-coil as observed for SS4. These helices are longer than the previous predicted cc region in SS5 (residues 68 to 120) (Abt et al., 2020).

The molecular model shows a dimeric organization of SS5, similar to that of SS4, involving two dimerization swapping-domains (from residue 121 to 139), one of each monomer (supplementary Figure 4C). The dimerization domain of SS4 is not conserved in SS5 sequence (Abt et al., 2020), however a dimerization domain is observed with a conserved structure when compared with SS4(supplementary Figure 4D). The helix H1 which interacts with the GT5 domain of the other monomer is involved in the predicted long N-terminal helices forming the coiled-coil region. It has been shown that when the previous predicted cc region of SS5 (residues 68-120) is absent, the protein is neither able to form dimers nor to interact with PII1 (Abt et al., 2020). This result is consistent with the molecular model of SS5, with the absence of the cc region likely to affect the structure of the dimerization domain.

The GT5 domain of SS5 is extended by a helix (439-460) positioned at the site occupied by the C-terminal helix originating from the GT1 domain of SS4 (which is absent in SS5) and interacting with the GT5 domain of SS5 (supplementary Figure 4E).

The model for SS5 is very similar, in terms of its N-terminal, cc, dimerization and GT5 domains, with the addition of the terminal helix, to SS4 at the dimerization site. The GT5 domain, the dimerization domain and the Helix 439-460 of SS5 superpose to the equivalent structures in SS4 with a rmsd of 1 Å on 250 Cα (supplementary Figure 4E). Only these two SSs have this dimeric organization and both interact with PII1, so it is conceivable that the dimerization region could be involved in the interaction with PII1.

### PII1 H2-H3 is soluble and organized in CC

To study PII1 and SS4, we made truncated constructs to obtain soluble samples compatible with biochemical and biophysical approaches. For PII1, we produced a protein construct containing only the H2-H3 helices that make up the conserved cc region, hereafter referred to as PII1-H2-H3.

PII1-H2-H3 is expressed in large amounts, most of which is found in the soluble fraction (Supplementary Figure5). The soluble fraction was incubated with IMAC beads and eluted with 500 mM imidazole. At this stage, and despite incubation with the beads, only part of the soluble sample was able to bind to the resin and eluted with imidazole. The fraction that did not bind was subjected to a second unsuccessful incubation with Ni-beads, suggesting that in solution the protein must adopt multiple conformations or aggregate and that in this form the 6-histidine tag is no longer accessible. The eluted fraction was then subjected to a steric exclusion chromatography (SEC) step. PII1-H2-H3 eluted in a very broad peak, probably due to its non-globular structure, and the presence of the long cc. This type of chromatography is probably not suitable for non-globular proteins and does not allow us to determine the monodispersity of the sample, but it did allow us to eliminate the contaminants present at the end of the affinity chromatography step. We therefore kept it in the protein purification protocol (Figure 4A).

**Figure 4:**
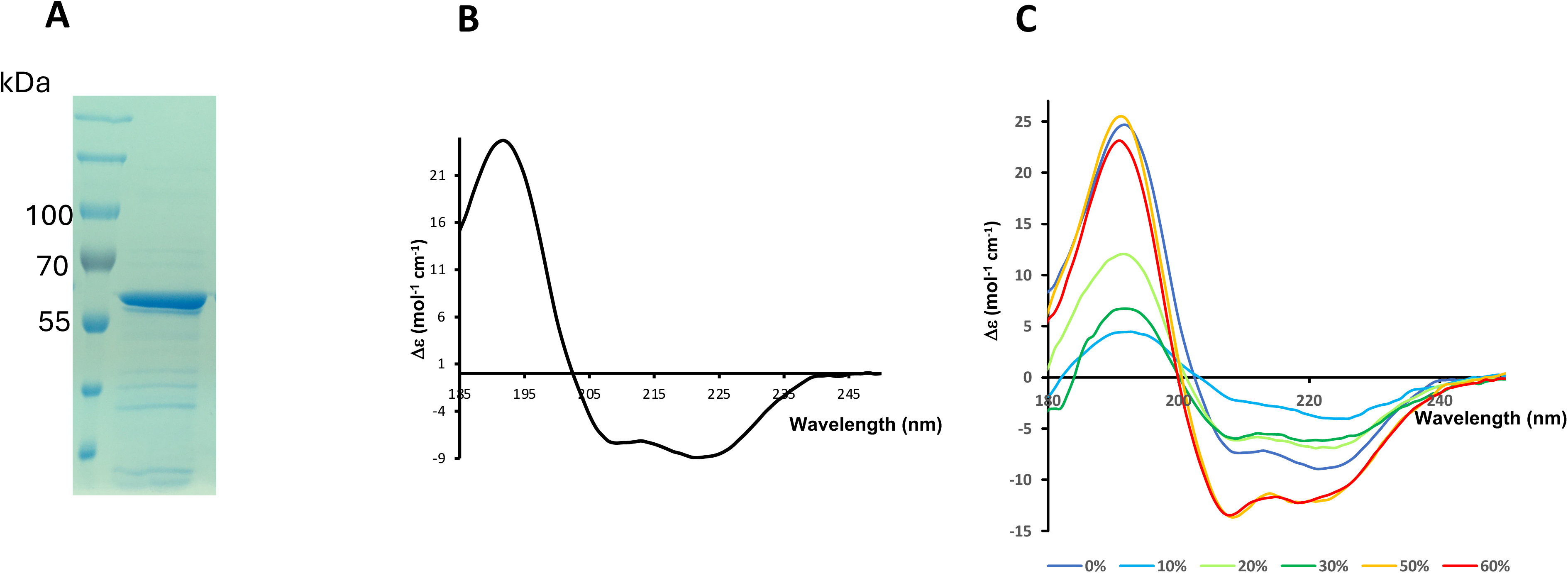
Purification and SR-CD analysis of PII1-H2-H3. **A)** SDS-PAGE of the protein sample of PII1-H2-H3 after purification, used for SR-CD experiments. **B)** SR-CD spectrum of PII1-H2-H3 **C)** SR-CD spectra of PII1-H2-H3 at different concentrations of TFE. The color code used for spectra in function of TFE concentration are indicated.

To experimentally verify the coiled-coil structure of PII1-H2-H3, we used a Synchrotron radiation circular dichroism (SR-CD) study approach on the purified PII1-H2-H3 protein sample at 3 mg/ml. This method is mostly used to analyze the content of secondary structure elements in a protein and to distinguish between individual helices and helices involved in a coiled-coil assembly, which is particularly relevant to our study. A typical spectrum obtained from alpha helices shows a positive peak around 190 nm and two negative peaks at 208 and 222 nm, the latter mainly due to helices, the others also coming from other structural elements. For free helices the magnitude of the 208 nm pic is lower than the one at 222 nm, the opposite is true for helices involved in a coiled-coil (Wallace, 2009). SR-CD analysis of PII1-H2-H3 presents a typical helix-folded protein profile (Figure 4B) with a positive peak at 190 nm and two negative peaks at 208 and 222 nm. The ratio of the ellipticity measured at 208 nm to that measured at 222 nm gives a value lower than 1, indicating and confirming the organization of the helices into cc.

To go further in the validation of the presence of cc regions, we measured different SR-CD spectra of PII1-H2-H3 in the presence of increasing amounts of 2,2,2-trifluoroethanol (TFE), which has the property of separating helices organized in cc and stabilize them as individual helices. The results show a rather atypical effect of TFE on PII1-H2-H3 at low concentrations of TFE (10%), the protein helices are initially unfolded, causing the spectrum to flatten. At 20-30% TFE, the helices gradually reform, resulting in a maximum signal corresponding to individualized helices at 50% TFE concentrations in which all helices are independent (Θ222/Θ208>1) (Figure 4C) thus definitely confirming the structuration of helices in cc in the structure of PII1. The fact that PII1 can easily unfold and then reform helices is quite atypical and shows that the protein is highly labile under certain conditions, allowing it to undergo major conformational changes.

### SS4-Δ349 is soluble and displays enzymatic activity

Since SS4 cannot be produced in its full form, we decided to express a truncated version of the protein for the biochemical assays. We based this on the work of (Raynaud et al., 2016), who expressed a soluble and active SS4 construct containing the cc3, cc4, dimerization and catalytic domains. The deleted part corresponds to the first 349 residues, so we named this construct SS4-Δ349.

SS4-Δ349 was expressed in small quantity but in soluble form. It was purified by an IMAC affinity chromatography step (Figure 5A).

**Figure 5:**
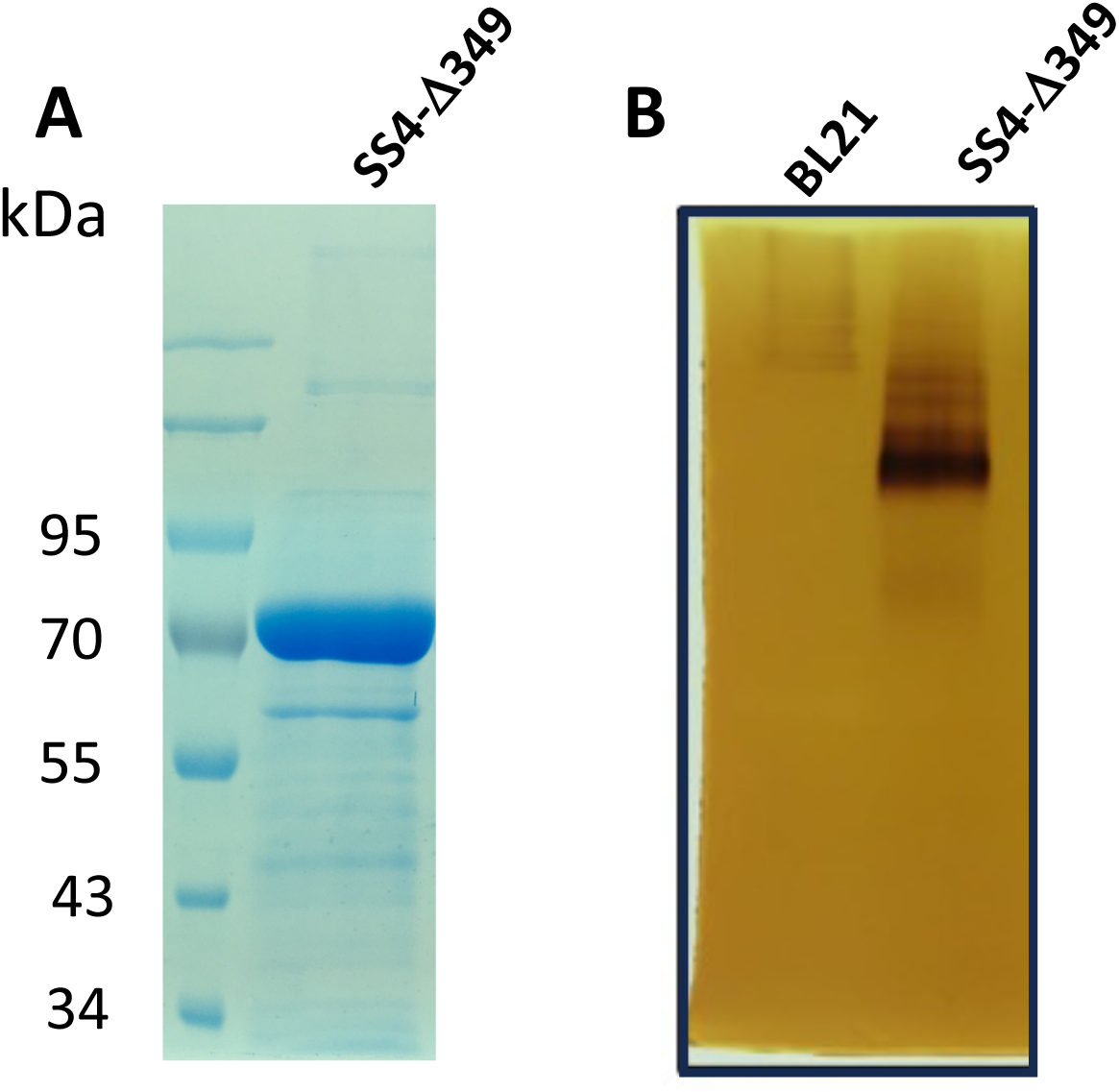
Purification and characterization of starch synthase elongation activity of SS4-Δ349. A) SDS-PAGE of the protein sample of SS4-Δ349 after purification B) Zymogram revealing the elongation function of SS4-Δ349. 150 µg of total soluble protein extract of BL21 (DE3) and BL21 (DE3) expressing SS4-Δ349 were loaded onto a 10 % non-denaturing acrylamide gel containing 0.3 % oyster glycogen (Sigma). After migration, the gel was incubated overnight at room temperature in synthase incubation buffer containing 2.4 mM ADP-glucose. Starch synthase activities were visualized as brown bands after iodine staining.

We then verified that deletion of cc1/cc2 in SS4 did not affect the ability of the enzyme to elongate glucan chains. To do this, we performed zymograms consisting of the separation by electrophoresis of soluble protein extracts of bacteria expressing SS4-Δ349 (150µg of soluble proteins), under native conditions, in a 10% polyacrylamide gel containing 0.3% glycogen. After migration we observed the capacity of the enzyme to elongate the outer chains of glycogen as revealed after iodine staining of the gels after incubation in the presence of ADP-Glc. This activity is revealed by the appearance of a brown/black coloring band with iodine, while the unmodified glycogen chains stains orange/light brown. To verify that the measured activity was not due to a bacterial enzyme, we performed the same experiment with a soluble extract of the same bacterial strain that does not express SS4-Δ349. (Figure 5B). Analysis of the results showed very low elongation activity for the BL21 strain likely corresponding to the *E. coli* glycogen synthase (GlgA) at the top of the gel. In contrast, a band of high elongation activity was observed in the strain expressing SS4-Δ349, indicating the high activity of the enzyme in its truncated form. This activity is shared by most of the starch synthases isoforms targeted to the chloroplast. It is clearly not this polyglucan elongation activity, even if it exists, that gives SS4 such an important role in plants, but its initiation activity. Studies using high performance anion exchange chromatography coupled to pulsed amperometric detection (HPAEC-PAD) showed that all SSs are capable of extending glucan chains from ADP-glucose and maltose or glucans with a higher degree of polymerization (DP) (Brust et al., 2013). None of them, including SS4, could synthesize *de novo* glucans from ADP-glucose and glucose (Brust et al., 2013). We tried to perform the same experiment by replacing glycogen with a mix of small MOS, but we were unable to detect any iodine staining.

### PII1 inhibits specifically SS4 activity

In order to check if the interaction with PII1 had an effect on the glucan elongating activity of SS4, we mixed soluble extracts from bacteria in which PII1-H2-H3 and SS4-Δ349 had been expressed. Since we could not measure the exact amount of each protein in the soluble extract, we first mixed an equivalent amount of total protein for each extract (175 µg and 350 µg, Figure 6A). After incubation for 15 minutes, the mixtures were loaded onto a polyacrylamide gel and the SS4 elongation activity was assessed as previously by iodine staining of the zymogram after incubation overnight with ADP-Glc (Figure 6A).

**Figure 6:**
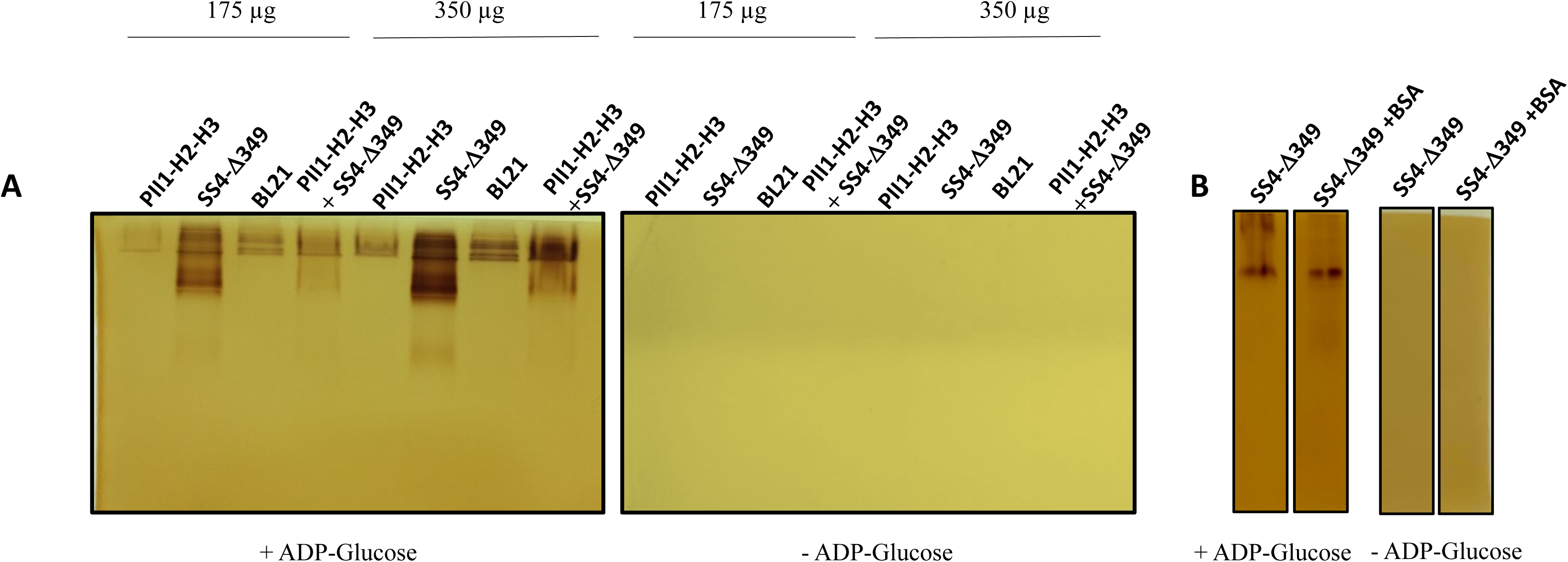
Zymogram revealing PII1-H2-H3 effect on SS4-Δ349 starch synthase activity. A) 175 or 350 µg of total soluble protein extract of BL21 (DE3) and BL21 (DE3) expressing SS4-Δ349 or PII1-H2-H3 were loaded onto a 10 % non-denaturing acrylamide gel containing 0.3 % oyster glycogen (Sigma). After migration, the gel on the left was incubated overnight at room temperature in synthase incubation buffer containing 2.4 mM ADP-glucose and the gel on the right in synthase incubation buffer without ADP-glucose. Starch synthase activities were visualized as brown bands after iodine staining. B) Zymogram analyzing the effect of BSA on SS4-Δ349 activity.

The ratio of each protein was estimated from the intensity of the PII1-H2H3 and SS4-Δ349 bands on SDS-PAGE. (Supplementary Figure 6A). This analysis showed that in the soluble extracts the amount of PII1 was about 60 times higher than that of SS4. This value should be treated with caution as the level of expression (especially for SS4) varies from one expression to another. For this experiment, we performed several controls in order to analyze the results unambiguously. First, we performed the same experiment without ADP-glucose to check that the observed activity was not linked to another substrate (Figure 6). We then performed the experiment with soluble proteins from expression bacteria that were not transformed by the plasmid to check that the observed activity was due to SS4-Δ349 and not to a bacterial protein (Figure 6A). We did the same with bacteria expressing PII1-H2-H3 to verify that this protein doesn’t have any enzymatic activity (Figure 6A). Finally, we checked that the activity of SS4-Δ349 remains unaffected in the presence of an excess amount of BSA (Figure 6B). To verify the specificity of the effect of PII1-H2-H3 on SS4-Δ349, we compared it with the effect on GlgA activity and tested the effect of increasing amounts of PII1 (Supplementary Figure 6B).

Analysis of the results shows that no elongating activity is observed in the absence of ADP-glucose, confirming that this precursor is used as a substrate by SS4-Δ349 and GlgA and that no effect of the presence of BSA could be observed on the activity of both SS4-Δ349 and GlgA (Figure 6B). PII1-H2-H3 doesn’t display any elongation activity.

In the presence of PII1-H2-H3, a strong inhibition of SS4-Δ349 activity is observed under the tested conditions (Figure 6A, Supplementary Figure 6B), and this inhibition is proportional to the amount of PII1-H2-H3 in the reaction mixture. However, the presence of PII1-H2-H3 does not affect the activity of bacterial GlgA as SS4-Δ349 loses its elongation activity whereas GlgA remains fully active regardless of the amount of PII1-H2-H3. This demonstrates the specificity of PII1-H2-H3’s inhibitory effect on SS4.

This experiment shows that PII1-H2-H3 interacts specifically to SS4-Δ349 and that this interaction inhibits SS4-Δ349 glucan elongation activity.

### Molecular modelling of the interaction between SS4 proteins involved in the initiation complex

Numerous interactions between the proteins involved in initiation have been demonstrated. PII1 can interact strongly with SS4, PTST2 and SS5 (Seung et al., 2018; Abt et al., 2020). PTST2 interacts strongly with PTST3 and MFP1 weaker with PII1 and transiently with the C-terminal domain of SS4 through its CBM (Seung 2017,2018). PTST3 interacts only with PTST2. To visualize the interaction modes of these proteins, we attempted to model all possible combinations of interaction using AlphaFold3 (Abramson et al., 2024). We calculated 50 structure models with AlphaFold3 for each complex and analyzed the results very carefully. Unfortunately, we were unable to obtain any models that met our selection criteria. In most cases we obtained models with reasonable pLDDT values, of the same order as the models obtained for the individual proteins, but the PAE matrices did not validate the positions of each molecule. Furthermore, of the 50 structure models calculated, most proposed different structures, sometimes even clashes between the different chains. The ipTM and pTM values never exceeded 0.35 and 0.4 respectively. This is probably due to several factors: most of the proteins involved in the initiation complex (all except the catalytic domain of SS4) have sections with very few homologous sequences, especially in the cc regions, regions with repeats which can be organized in a large number of structural variations, which makes the training and subsequent inference harder, especially when several partners are involved, knowing the variety of binding they can be involved in (see discussion), and all the proteins have disordered regions which reduce the prediction scores of the obtained models

## Discussion

All the proteins involved in starch binding initiation have been proposed to interact with each other to form an initiation complex (Seung et al., 2018; Abt et al., 2020). This proposal was supported by interaction studies between the identified molecules and by the fact that all these proteins have more or less important regions of their structure predicted to be organized in coiled-coils, known to be involved in protein-protein interactions (Truebestein and Leonard, 2016).

As their physical properties, particularly their length and flexibility, have important structural and functional properties, the cc are widely found in biological processes and multimolecular complexes (for review see (Truebestein and Leonard, 2016). Long cc, act often as molecular spacers, either separating functional domains or scaffolding large macromolecular complexes. Some cc act simply as molecular spacers or facilitating oligomerization, while others have evolved the ability to communicate conformational changes along their length. Some of them are able to bind membranes, link two compartments in the cell (Engel et al., 1992) conferring conformational plasticity. The coiled-coil domain of myosin, which is similar to PII1, has been shown to act as a mechanosensor and to transmit structural changes along its chain (Linari et al., 2015).

At the present stage of our knowledge of initiation mechanisms, we know that all the proteins involved have one or more cc regions predicted on the basis of sequence, and we have information on interactions between proteins that have been shown experimentally. However, apart from the crystallographic structure of the catalytic domain of SS4 and the CBM48 domains of the PTSTs, there is no structural information and very little knowledge about the regions involved in the interactions. Nor do we know whether the initiation complex involves all the proteins at once, or whether the interactions are transient or sequential.

In the present work we focused on the structural study of PII1 and SS4 combined with structural and biochemical approaches to determine the effect of the binding of PII1 on SS4 enzymatic activity.

### PII1 contains a long conserved coiled-coil domain

We studied the structure of PII1 using molecular modelling and showed that the protein is composed of 6 helices that are conserved in all identified orthologs found on the TAIR website except *Setaria italica*, which has a shorter sequence corresponding only to the H1 H2 and H3 helices of *At*PII1. The conserved sequences also adopt a conserved 3D structure, in particular the long H2-H3 helices, which account for almost 60% of the amino acids in the protein. These two helices join together to form a long cc region, which is about 300-350 Å in length. The presence of cc was demonstrated experimentally by SR-CD, but could not be observed by SAXS due to the conformational heterogeneity of the solution. We have shown that this domain is conserved in all tested orthologs and that it may play an important role in the function of PII1.

At this stage, it is difficult to envisage a function for PII1 solely on the basis of its structure. However, it has been shown that PII1 interacts with PTST2, SS5 and SS4, and it may be involved in regulating SS4 activity.

### What is the advantage of organizing SS4 and SS5 into dimers?

Both SS4 and SS5 form dimers (Raynaud et al., 2016; Abt et al., 2020). We showed that a structurally conserved dimerization region located between the N-terminal cc region and the C-terminal catalytic domain is involved in the dimer formation. Our results show with high confidence that these dimerization regions form a small globular domain that interacts with the GT5 subunit of the SS4 catalytic domain or the SS5 C-terminal domain.

This organization, in which each monomer exchanges its dimerization domain, interacting strongly with the GT5 domain of the second monomer, offers some interesting properties. Firstly, it allows the two GT5 domains of the catalytic domains to be kept close together, leaving the catalytic region (for SS4) or the glucan-binding domain (for SS5) accessible. On the other hand, the dimerization domain forms a hinge region between the GT5 domains and the cc regions, the structure of which confers dynamic properties that allow signals to be transmitted from the cc domain to the catalytic domain *via* conformational changes induced by the binding of other partners. It is therefore conceivable that the binding of PTST2 and/or PII1 to SS4 could influence its catalytic activity in the initiation complex.

SS4 and SS5 both interact specifically with PII1 which does not recognize other families of SSs.

All SSs share a catalytic domain (for functional SSs) or at least the GT5 domain and non-conserved N-terminus domains. SS1 and SS2 have no predicted cc regions, in contrast to SS3, SS4 and SS5 (Abt et al., 2020). Only SS4 and SS5 interact with PII1. The specificity of their recognition by PII1 cannot be entirely related to the presence of cc regions, since SS3 is not recognized. Nor can it be due to the structure of GT5, which is present in all SSs. What distinguishes SS4 and SS5 from other SSs is the dimerization associated with the presence of a structurally conserved dimerization domain that interacts strongly with the GT5 domain and acts as a hinge with the cc region. Since this dimerization is necessary for recognition by PII1 (Abt et al., 2020), it is conceivable that the latter interacts with SS4 and SS5 specifically with this part of the molecule and (due to the structure of PII1) with the coiled-coil regions of the SSs. Dimerization could therefore be the prerogative of SSs involved in the initiation of starch biosynthesis. This also implies that PII1 interacts with either SS4 or SS5, but not both at the same time. In this case, SS5 could have a regulatory effect on the binding of PII1 to SS4.

### PII1 can inhibit SS4 enzymatic activity

Apart from the fact that it interacts with SS4, PTST2 and SS5 and that its absence in the plant results in a phenotype in which the initiation of starch granule synthesis is affected, the role of PII1 is not known. In this paper, we have shown that PII1 conserved cc interacts with a truncated form of SS4 and that this interaction inhibits the elongating activity of SS4. We have also shown that this inhibition is specific to SS4 and does not affect the activity of the *E.coli* glycogen synthase GlgA. Since GlgA and SS4 have similar catalytic domains, this result indicates that PII1 probably does not interact directly with the catalytic domain of SS4, but rather *via* the cc domain of the enzyme (which is also part of the dimerization) and/or its dimerization region.

It is rather counterintuitive to observe the inhibition of SS4 by PII1, since the absence of PII1 in the plant leads to a reduction in initiation events. This inhibition can only occur transiently in a complex, highly dynamic mechanism in which interactions within the initiation complex are transient or evolving. We have therefore re-examined the current data in the light of our results.

As shown by the structural analysis, the dimerization domain of SS4 forms a hinge region between the cc region and the catalytic domain. It is likely that binding of PII1 to SS4 induces a conformational change in the catalytic domain leading to its loss of activity maybe braking the link between the two catalytic domains or separating both GT domain. However, the strength of this inhibition must be qualified in the absence of other initiation proteins, in particular PTST2, which binds to the catalytic domain of SS4 *via* its CBM48 domain (Seung et al., 2017). PTST2 interacts as well with MFP1 and this interaction is involved in the localization of SS4 to thylakoid membranes (Sharma et al., 2024). The sites of interaction between PTST2, PII1 and MFP1 are not known, but since the latter two are almost entirely cc-structured, it is likely that this interaction occurs *via* the cc region of PTST2. As PII1 has conserved helices (outside the cc domain) and PII1 and PTST2 have predicted structurated and unstructured regions, we cannot rule out that these regions are involved in the interaction, increasing the possibility of signal transduction by conformational change. Our circular dichroism results showed that PII1 could easily reorganize into other structures, indicating a certain lability in its structuring.

The CBM48 domain of PTST2 binds specifically helically folded glucans with DP 10 or higher that are the shortest MOS able to form crystalline glucans. It has been proposed that PTST2 brings these substrates to SS4 to induce initiation (Seung et al., 2017). However, in a yeast model system expressing different combinations of starch biosynthetic enzymes, the presence of SS4 tended to promote the formation of insoluble glucans, and this effect was seen in the absence of PTST2 (Pfister and Zeeman, 2016). These glucans can be used as substrate by SS1 for amylopectin biosynthesis. If such substrates exist in the chloroplast, why would an initiation step be required if SS1 can use them? On the other hand, PTST2 is required for the initiation of biosynthesis and is also responsible for poly-initiation when overexpressed in plants. Another hypothesis for its function may be that helical glucans would not be substrates but products of the reaction catalyzed by SS4. The function of PTST2 may be to receive these products and make them available to the enzymes involved in starch biosynthesis. In that way PII1, which inhibits SS4 activity, could intervene to stop the elongation of glucan chains once they are sufficiently long to emerge from the catalytic region, at which point they would be retrieved by PTST2. Once released, the polyglucan could be taken up by the enzymatic biosynthetic machinery and PTST2 could reposition itself to suppress inhibition by PII1 allowing then SS4 to initiate a new primer. Our hypothesis is that in the absence of PII1, SS4 initiates the synthesis of the glucan chain, but since the enzyme can no longer be inhibited by PII1, it behaves like the other SSs and forms much longer chains, reducing the number of primers available and therefore the number of initiation events. SS5 may play a role in regulating PII1 function.

In conclusion, the fact that each of these molecules has one or more cc regions, giving them the ability to interact with each other, but also to transmit conformational change signals within the initiation complex itself, suggests a complex and highly regulated mechanism of which we currently know only the basics. Our current knowledge is based on analyses of the phenotypes of mutants in which the initiation proteins have been inactivated or overexpressed, and on identification of partners, which shed light on the nature of the interactions but do not allow us to go any further. In this work, we have studied the structural and enzymatic aspects of the current state of knowledge, which has allowed us to make progress in our understanding of the mechanisms and also to put forward some new hypotheses. The difficulty of producing the key players in the initiation complex in solution, and the presence of long cc regions that tend to aggregate in the absence of partners, complicates conventional structural biology tools that require monodisperse solutions. Determining the stoichiometry of each complex formed in solution and using cryo-electron-microscopy to study them will undoubtedly be an asset in the study of these complex mechanisms. The development of AI tools for modelling molecular models of multi-protein complexes, which is rapidly expanding, should soon make it possible to predict the interactions between the different initiation proteins and greatly improve our understanding of these mechanisms.

## Acknowledgements and funding

The authors strongly acknowledge DISCO beamline at synchrotron SOLEIL (St Aubin, France) and are grateful for the expert technical support provided by beamline staffs. The authors would like to thank Dr. Fabrice Wattebled for providing plasmids expressing full-length PII1 and SS4 and helpful discussions. This work has been supported by Centre National de la Recherche Scientifique France (cora funding to G.B, M.L., D.D., C.B. and PhD grant for R.O.), Université de Lille France (PhD grant to M.B), and Région Hauts de France (PhD grant for M.B.).

## Author Contributions

CB, MB and DD conceived and designed the experiments. MB performed experiments with input from CB, RO, DD, ML and GB. CB, RO and MB collected synchrotron data. MB and CB analyzed data with the input of DD. CB wrote the manuscript. CB, MB DD, GB and ML revised the manuscript.

---

**Supplementary Figure 1:**
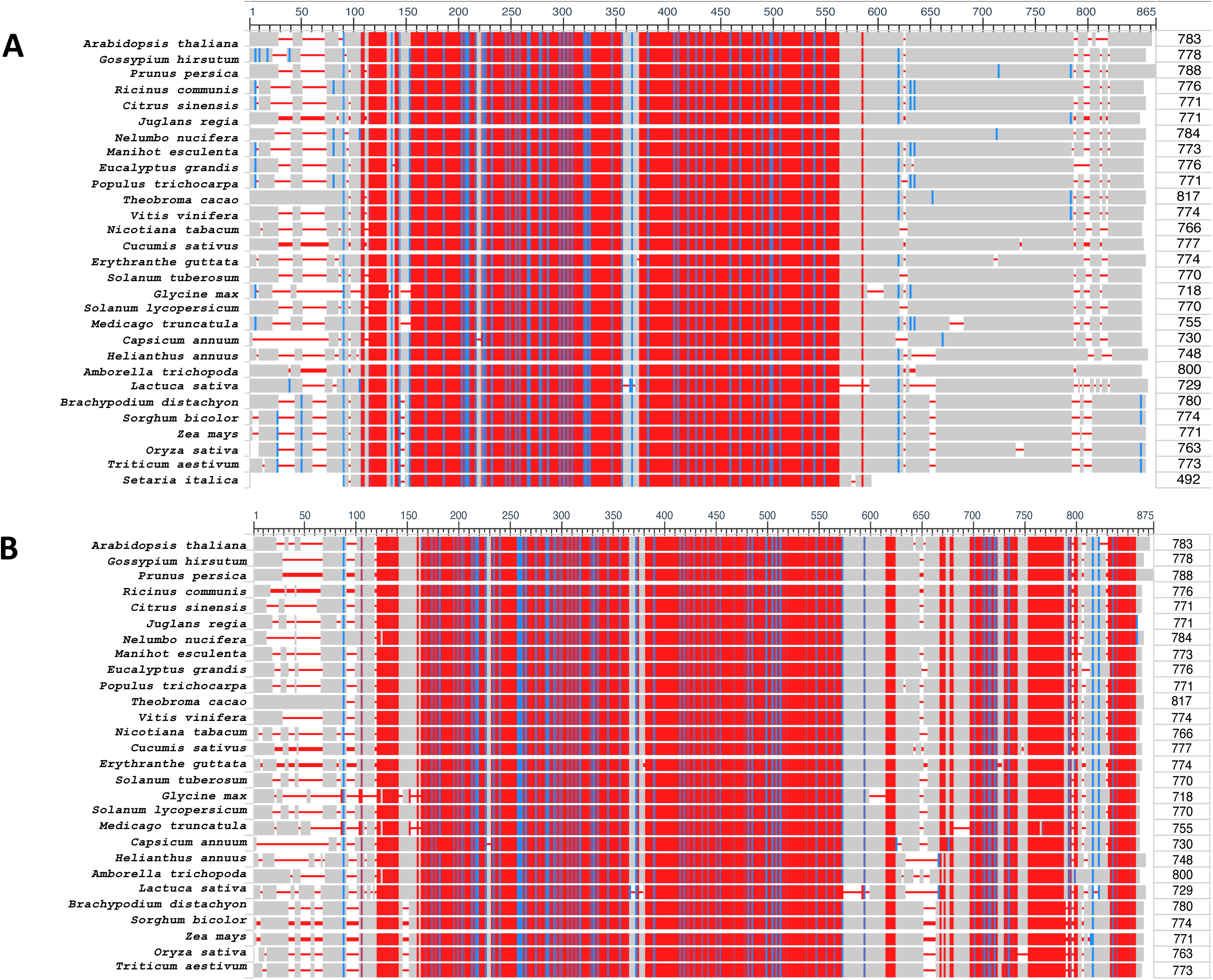
Primary structure alignment of *At*Pii1 with plant orthologs. A) Sequence alignment of *At*PII1 with all the analyzed orthologs B) sequence alignment of *At*PII1 orthologs except ortholog from *Seratia italica*

**Supplementary Figure 2:**
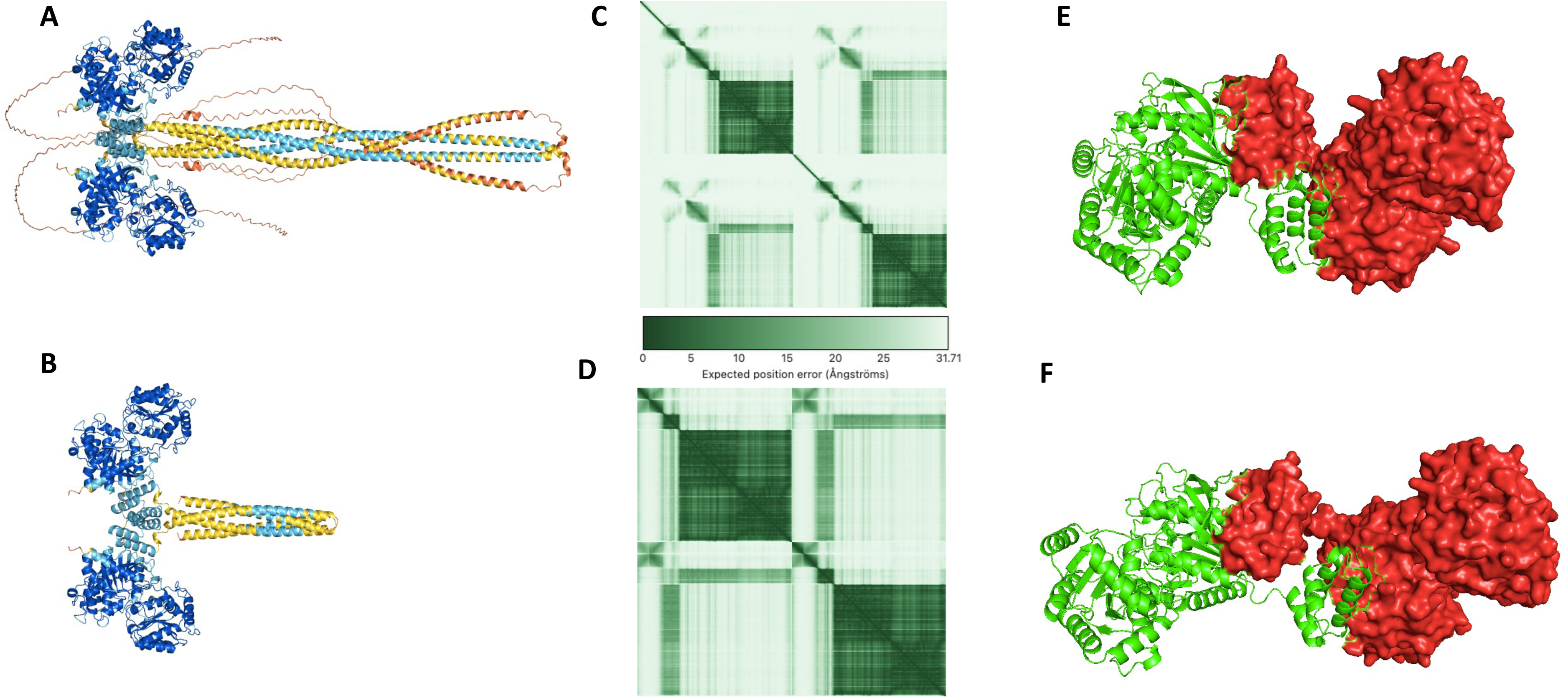
AlphaFold models of dimeric SS4 and SS4-Δ349. A) structure of SS4 B) and SS4-Δ349. regions with pLDDT > 90 are colored navy blue, regions with 90 >pLDDT > 70 are colored cyan, regions with 70 >pLDDT > are colored yellow and 50 pLDDT < 50 are colored orange. C) PAE matrix for SS4 D) and SS4-Δ349. E) Dimeric organization of SS4 and F) SS4-Δ349. Only the catalytic and dimerization domains of each monomer are represented. One monomer is represented as green cartoon and the second is represented as red molecular surface.

**Supplementary Figure 3:**
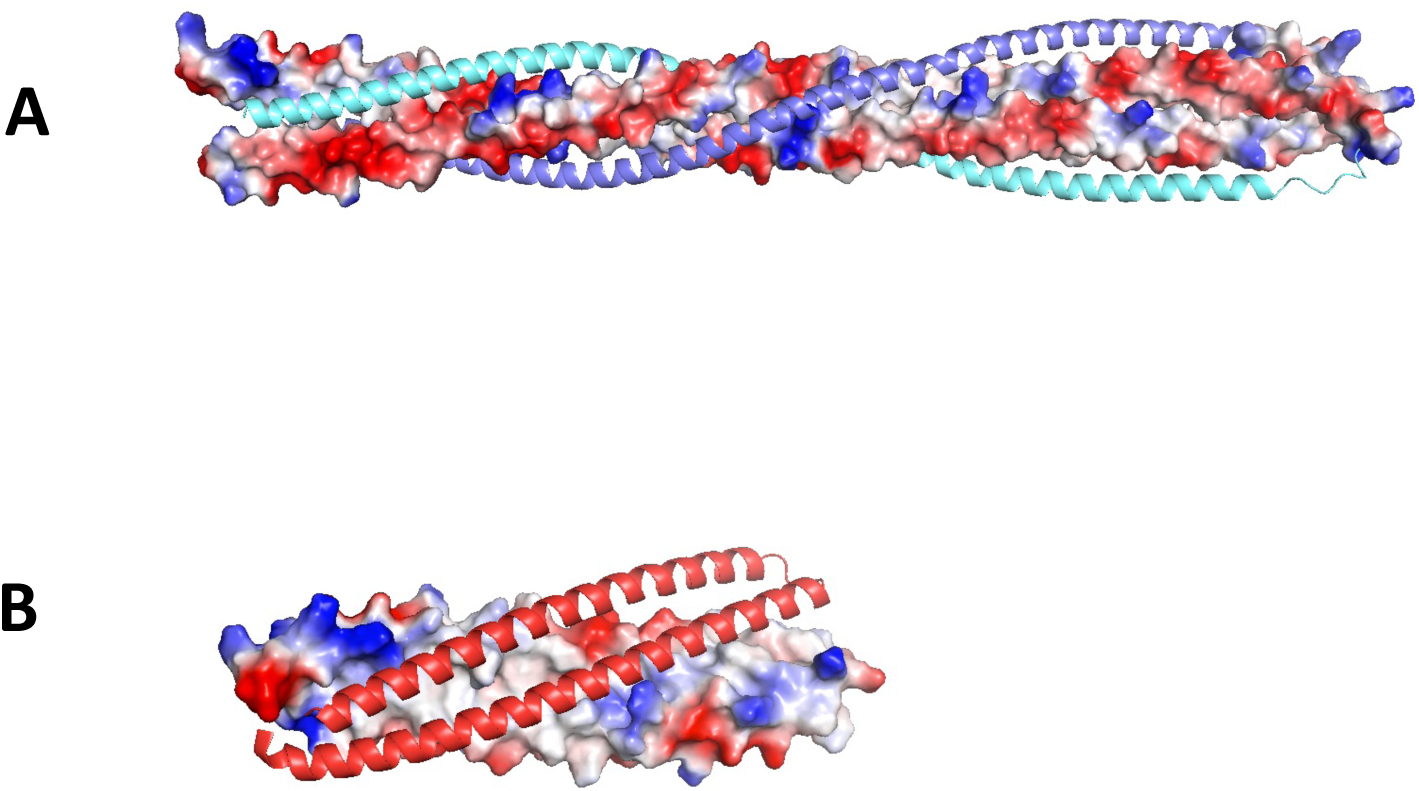
Structural analysis of the N-terminal tetracoil regions of SS4 and SS4-Δ349 dimers. A) In SS4, the two cc3/cc4 helices forming a central dimeric coiled-coil are represented as surface potential and the cc1-cc2 helices that are supercoiled around are represented as cartoon. B) In SS4-Δ349 dimers, the cc3 and cc4 regions form distinct helices forming an antiparallel cc that assembles with the cc3/cc4 region of the second molecule. The cc3 and CC4 helices of one monomer are represented in surface potential whereas the cc3 and cc4 helices of the second monomer are represented in cartoon.

**Supplementary Figure 4:**
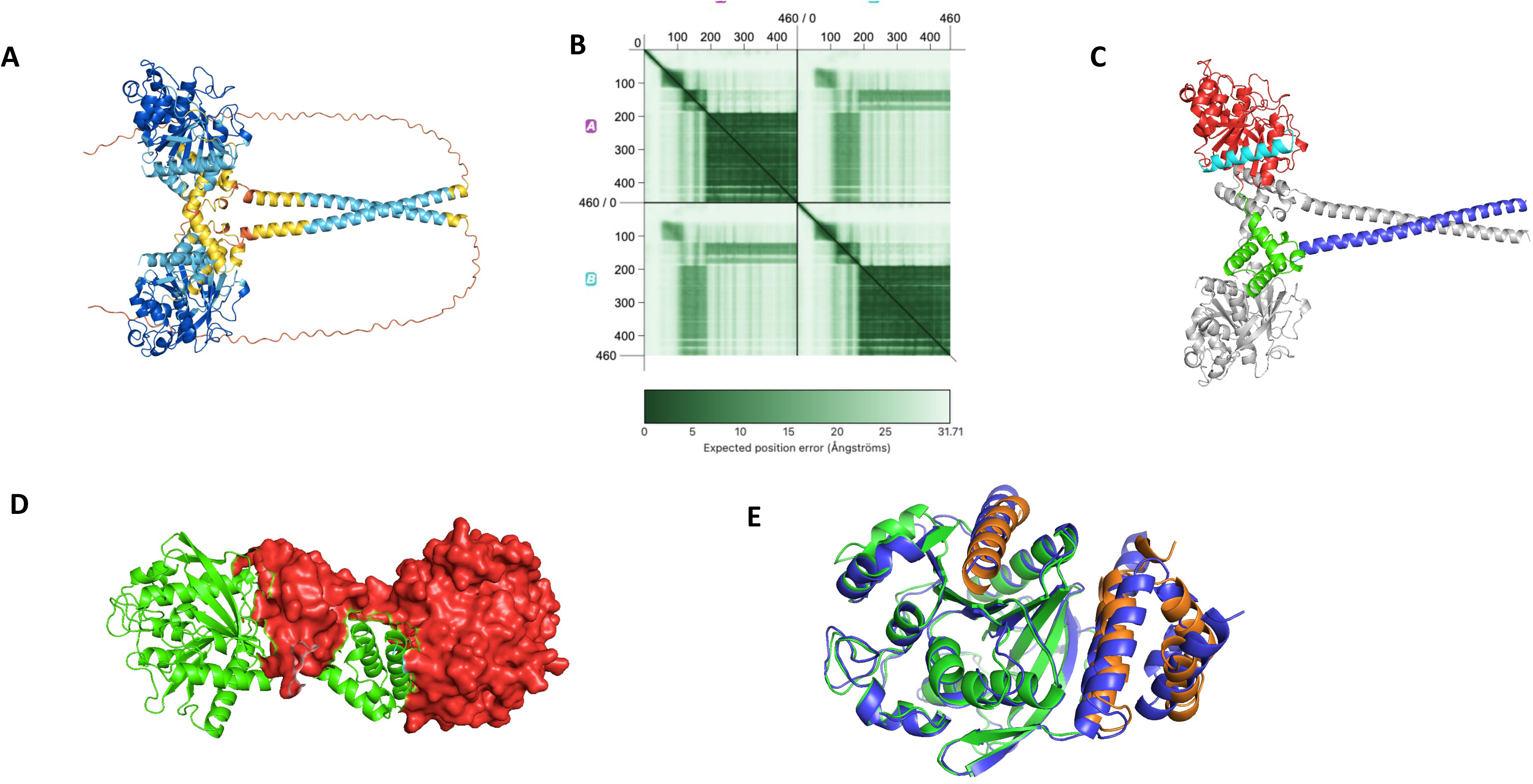
AlphaFold models of SS5 dimers. A) Structure of SS5, regions with pLDDT > 90 are colored navy blue, regions with 90 >pLDDT > 70 are colored cyan, regions with 70 >pLDDT > are colored yellow and 50 pLDDT < 50 are colored orange. **B)** PAE matrix for SS5 dimer **C)** structural organization of SS5. Only one monomer is colored (GT5 domain in red, dimerization domain in green, the helix (439-460 in cyan). The second monomer is colored gray **D)** Dimeric organization of SS5. Only the GT5 and dimerization domains of each monomer are represented. One monomer is represented as green cartoon and the second is represented as red molecular surface. **E)** Structure superposition of the GT5 and dimerization domains of SS5 (in purple) and SS4 (in green for GT5 and orange for the dimerization domain and the C-terminal helix of GT1).

**Supplementary Figure 5:**
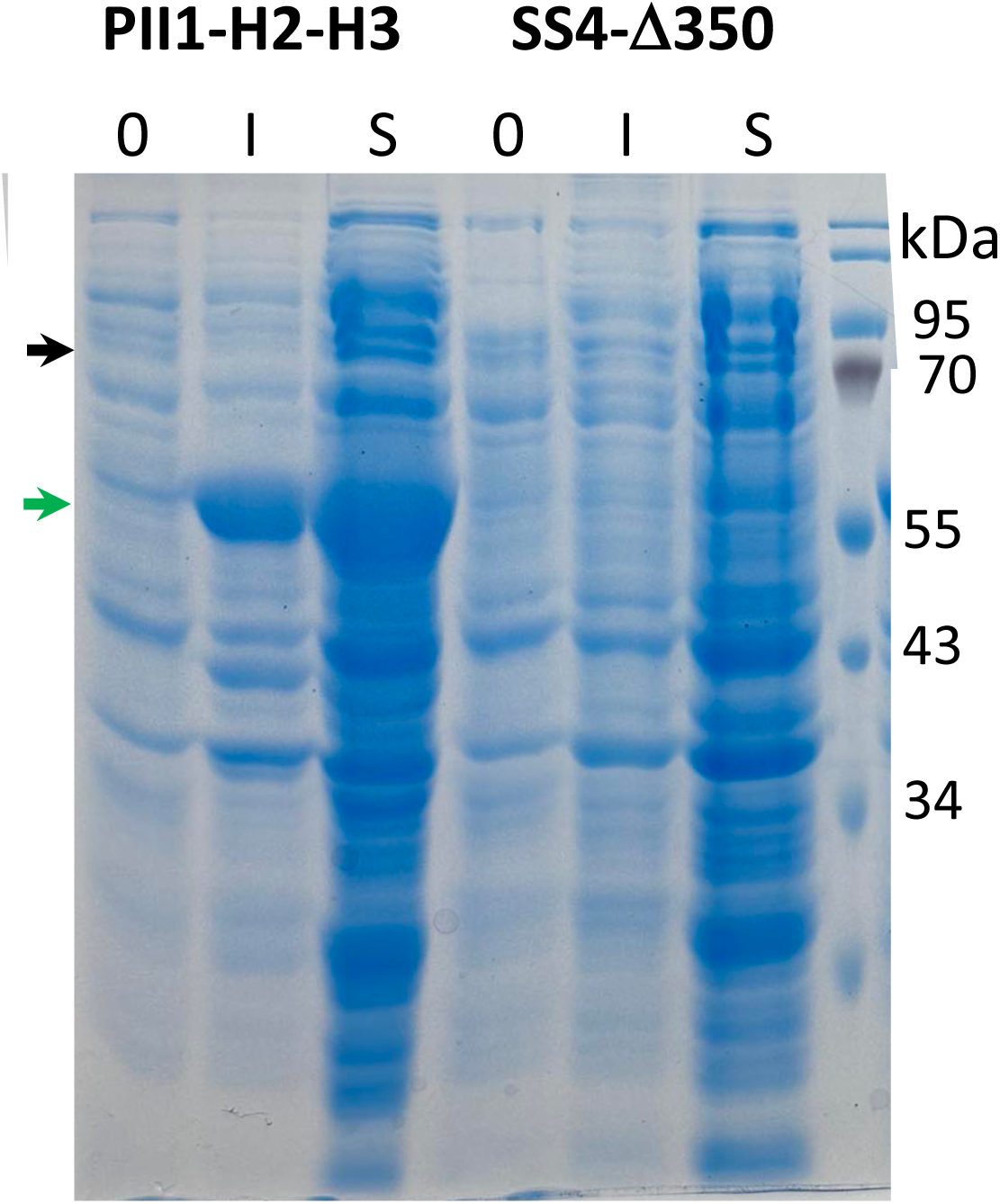
SDS-PAGE showing expression and solubility of PII1-H2-H3 and SS4-Δ349. Crude bacterial extracts before induction (0), after induction (I) and soluble fractions (S) have been loaded on a 10% SDS-PAGE. Migration bands of PII1-H2-H3 and SS4-Δ349 are indicated by green and black arrows respectively.

**Supplementary Figure 6:**
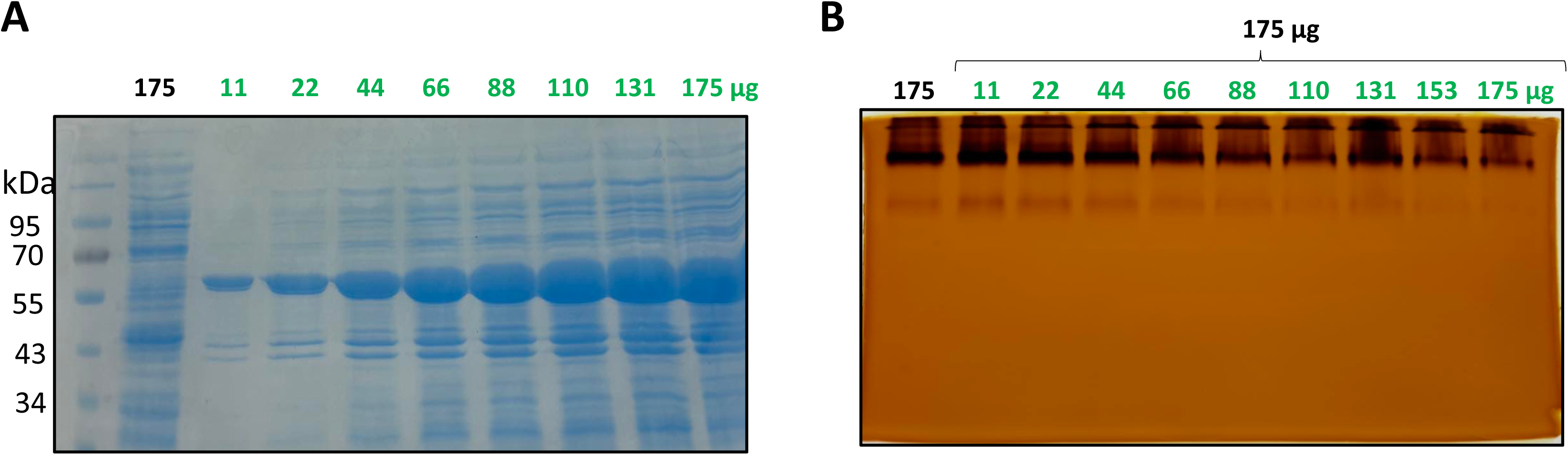
Influence of SS4-Δ350 / PII1-H2-H3 ratio on SS4 elongation activity. A) SDS-PAGE loaded with different quantities (in µg) of SS4-Δ350 (in black) and PII1-H2-H3. (in green) bacterial soluble extract. B) Zymogram revealing the effect of increasing amounts of PII1-H2-H3 on SS4-Δ349 starch synthase activity.

